# Evolutionary models predict potential mechanisms of escape from mutational meltdown

**DOI:** 10.1101/2022.06.21.496937

**Authors:** Claudia Bank, Mark A. Schmitz, Ana Yansi Morales-Arce

## Abstract

Mutagenic drugs are promising candidates for the treatment of various RNA virus infections. Increasing the mutation rate of the virus leads to rapid accumulation of deleterious mutation load, which is proposed to ultimately result in extinction as described by the theoretical concepts of mutational meltdown and lethal mutagenesis. However, the conditions and potential mechanisms of viral escape from the effects of mutagenic drugs have not been conceptually explored. Here we apply a computational approach to quantify the population dynamics and genetics of a population under high mutation rates and discuss the likelihood of adaptation to a mutagenic drug by means of three proposed mechanisms: (1) a proportion of “traditional” beneficial mutations that increase growth/fitness, (2) a mutation rate modifier (i.e., evolution of resistance to the mutagenic drug) that reduces the mutation rate, and (3) a modifier of the distribution of fitness effects, which either decreases or increases deleterious effects of mutations (i.e., evolution of tolerance to the mutagenic drug). We track the population dynamics and genetics of evolving populations and find that successful adaptations have to appear early to override the increasing mutational load and rescue the population from its imminent extinction. We highlight that the observed stochasticity of adaptation, especially by means of modifiers of the distribution of fitness effects, is difficult to capture in experimental trials, which may leave potential dangers of the use of mutagenic treatments unexposed.

## 1 INTRODUCTION

The search for therapeutics against virus infections remains a challenge in human health research. Understanding within-host infections and the potential for adaptation (including resistance) to new drugs requires studying the eco-evolutionary dynamics of virus populations. One heatedly discussed potential treatment option is particularly rooted in evolutionary theory: mutagenic drugs like favipiravir or molnupiravir act by inducing mutational meltdown (1, 2, 3) or lethal mutagenesis (4, 5); for a discussion of the respective terms see Matuszewski et al. (6). Mutational meltdown occurs when the virus population accumulates so many deleterious mutations that the population growth rate falls below 1, which induces extinction. Importantly, the evolutionary theory of mutational meltdown rests on the knowledge that most new mutations are deleterious. Moreover, the action of a mutagenic drug is a population process. Other drugs act by killing individual cells or viruses, or by stopping their proliferation. Conversely, a key aspect of mutational meltdown, induced by a mutagenic drug, is that the population as a whole is weakened through the successive accumulation of deleterious mutations, which ultimately leads to extinction.

Mutagenic treatment is particularly promising against RNA viruses because of their comparably high mutation rates (7, 8), which suggests that only a slight increase in the mutation rate is sufficient to induce mutational meltdown (9, 10). Mutagenic drugs such as favipiravir and molnupiravir increase the mutation rate by targeting the RNA polymerase. After phosphorylation inside the cell their active forms become incorporated as nucleoside analogues in the newly synthesized RNA chain (11, 12, 13, 14). Experimental and clinical trials of mutagenic drugs against Sars-Cov-2 have shown promising results (15, 16, 17), although Sars-Cov-2 has a relatively low mutation rate as compared to other RNA viruses. However, recent clinical data indicated that their effectiveness was inversely proportional to the severity of symptoms (18) or completely absent (19). A possible explanation is a counteracting proof reading mechanism called ExonN, that avoids the incorporation of mutagenic drugs into the virus genome (20, 21). The combination of different mutagenic drugs, able to overcome a proofreader, to treat Covid-19 patients seems still promising and is currently under debate (22, 23).

Several examples of adaptation of RNA viruses to mutagenic drugs have been reported (reviewed in (24). For example, rivabirin resistance was observed *in-vitro* in poliovirus and patient hepatitis C virus strains (25, 26), whereas remdesivir was shown to induce resistance in *in-vitro* Sars-CoV-2 (27). High doses of the mutagenic drug favipiravir did not help patients that were infected with the Ebola virus (28) and resistance was also described in chikungunya virus and in enteroviruses (29, 30). Yet, favipiravir is one of the most effective compounds inducing extinction in *in-vitro* influenza A virus, foot-and-mouth disease virus, and West Nile virus (13, 31, 32). Across various screens in the laboratory in influenza virus (12, 33, 34, 35), only one potential resistance mutant against favipiravir has been identified and experimentally validated to date (34, 35). Because of its special mode of action, it was hypothesized that the effect of mutagenic drugs may depend on the viral replication system, the genome structure, and the time at which treatment starts (10, 23).

Although the foundation for the proposed success of mutagenic treatments lies in evolutionary theory, evolutionary theory of mutational meltdown also predicts that mutagenic drugs are not without risks (36). Although most new mutations are deleterious, a mutagenic drug increases not only the chances of new deleterious but also of new beneficial mutations. Increasing the mutation rate could allow the virus population to “find” fit genotypes (e.g., including multiple, individually deleterious mutations) that would be unreachable under the default mutation rate. The longer a virus population survives under mutagenic drug treatment, the larger the possibility that adaptive genotypes may appear, which could cause damage not just in the current host, but also when transmitted to new, untreated, hosts (37, 38).

Here we approach this problem by proposing three mechanisms of adaptation to mutagenic drugs from the view of an evolutionary biologist. Using computer simulations, we quantify the eco-evolutionary dynamics of a clonal (virus) population under high mutation rates that has access to either of these mechanisms and show that each can lead to escape from mutational meltdown under different conditions. Because the actual eco-evolutionary dynamics of a virus population inside a human host are complex and many of the parameters are unknown, we restrict ourselves to a small and fixed set of parameters. Thus, our study is meant as a proof-of-concept to spur discussion and future work.

We discuss the escape of a virus population from mutational meltdown through three adaptation mechanisms:

1. Beneficial, growth-rate increasing mutations. Because accumulation of deleterious mutation load is unstoppable by such mutations, a continual input of a fraction of growth rate increasing mutations is necessary to save the population from extinction that strongly depends on the carrying capacity.
2. A mutation rate modifier, which is a mutation that reduces the mutation rate of the virus genome-wide. We show that this type of mutation has to invade early to lead to escape from mutational meltdown, and argue that this mechanism qualifies as evolution of *drug resistance*.
3. A modifier of the distribution of fitness effects (in short: DFE modifier), which is a mutation that either dampens or exaggerates the deleterious effect of all mutations that its carrier is accumulating. We show that both dampening and exaggerating mutational effects can lead to escape from mutational meltdown and argue that this mechanism qualifies as evolution of *drug tolerance*.

The distinction between resistance and tolerance comes from the field of host-pathogen interactions (39, 40). We have not seen these concepts discussed in the evolutionary literature with respect to different mechanisms of adaptation. We show below that adaptation via evolution of resistance *versus* tolerance lead to fundamentally different implications for the eco-evolutionary dynamics of virus populations in the presence (and absence) of a mutagenic drug.

## 2 MATERIALS AND METHODS

### 2.1 Eco-evolutionary dynamics without adaptation

Similar to Lansch-Justen et al. (41), we model the intra-host eco-evolutionary dynamics of a non-recombining asexual virus population with a carrying capacity of *K*, which expands from a founding population size of *N*_0_ mutation-free individuals. The initial absolute growth rate (i.e., the average number of offspring that a mutation-free genotype produces) is *w*_0_. Each generation, every individual gains a number of deleterious mutations drawn from a Poisson distribution with mean *U*_del_. We consider an infinite-site model, i.e., every mutation occurs at a new site in the genome; back mutations are neglected. After mutation, the offspring number, i.e. the number of clonal copies of each individual, is drawn from a Poisson distribution with its mean set to the absolute growth rate *w*(*k*) of the respective individual, and the individual subsequently dies. We assume that each deleterious mutation has the same effect *s*_del_. We consider a multiplicative model that determines the absolute growth rate of an individual, *w*(*k*) = *w*_0_ *·* (1 + *s*_del_)^*k*^, where *k* is the number of mutations that the individual has accumulated. Finally, if the population size is above the carrying capacity *K*, individuals are eliminated randomly to adjust the population size to *K*.

### 2.2 Modeling the three adaptation mechanisms

We separately study three different models of adaptation mechanisms. In Section 3.2 we study growth-rate increasing mutations that occur at mutation rate *U*_ben_ with effect size *−s*_del_. Here, the absolute growth rate of an individual is *w*(*k, i*) = *w*_0_ *·* (1 + *s*_del_)^*k*^(1 *− s*_del_)^*i*^, where *i* denotes the number of beneficial mutations the individual carries. Thus, every beneficial mutation neutralizes one deleterious mutation.

In Section 3.3 we study mutation rate modifiers. A modifier of the mutation rate occurs at rate *U*_mutMod_ per genome per generation. The absolute growth rate of individuals remains the same as in the basic model, but the mutation rate of carriers of at least one mutation modifier is reduced to *U*_del_ *· f*_mutMod_, where *f*_mutMod_ is the effect size of the mutation modifier. The effect size is defined multiplicatively, i.e., if *f*_mutMod_ = 1, the mutation rate remains unaltered by the modifier mutation, and if *f*_mutMod_ = 0.1, the mutation rate is reduced to 10% of its original value.

In Section 3.4 we study modifiers of the distribution of fitness effects (DFE). A DFE modifier occurs at rate *U*_dfeMod_ per genome per generation. Here, the absolute growth rate of a carrier of at least one DFE modifier mutation is *w*^*∗*^(*k*) = *w*_0_ *·* (1 + *s*_del_ *· f*_dfeMod_)^*k*^, where *f*_dfeMod_ is the effect size of the DFE modifier. Thus, if *f*_dfeMod_ = 1, the modifier mutation leaves mutational effects unaltered, whereas DFE modifiers with *f*_dfeMod_ *>* 1 make every existing and new mutation more deleterious, and DFE modifiers with *f*_dfeMod_ *<* 1 make every existing and new mutation less deleterious.

### 2.3 Simulations and choice of parameters

Throughout the paper, we show results from simulations of the above-described models with carrying capacities *K* ∈ {500, 10000} and a founding population size *N*_0_ = 10. Each simulation ends when the population goes extinct, or after 5000 generations. We set the initial absolute growth rate to *w*_0_ = 2 and the deleterious mutation rate per genome per generation to *U*_del_ = 0.2. In Sections 3.2-3.4, the effect size of a deleterious mutation is *s*_del_ = *−* 0.023. In the absence of any adaptation mechanism, the mean extinction time for our standard parameter set for *K* = 500 is 592.44 *±* 69.28. Here, the minimum and maximum extinction times observed in *>* 10000 simulations were 384 and 931, respectively. For our standard parameter set for *K* = 10000, the mean extinction time is 1377.46 *±* 157.36. The minimum and maximum extinction times observed over *>* 10000 simulations were 947 and 2141 respectively. All results are based on simulations implemented in Julia v.1.0 or v.1.6.0. The annotated simulation and analysis code is available at https://gitlab.com/evoldynamics/ms-mutational-meltdown-2021 and will be archived on Zenodo upon publication of the manuscript.

## 3 RESULTS

### 3.1 Known features of extinction under mutational meltdown

Since the 1990’s, many theoretical studies have quantified mutational meltdown (41). We here reiterate some of these results using our simulations to familiarize the reader with the most important features of this evolutionary process. As described in Lynch et al. (3), Lansch-Justen et al. (41), the path to extinction by mutational meltdown consists of three phases. In the first phase, a clonal, non-recombining, deleterious-mutation-free population invades a new host, where it expands and accumulates deleterious mutations. Eventually, due to the combined effects of random genetic drift and mutation pressure, the fittest genotype will be lost, which marks the beginning of the second phase. In the second phase, Muller’s ratchet leads to the step-wise loss of the fittest genotype and an almost linear decrease in mean fitness, until the absolute growth rate of the population falls below 1. At this point, the third phase of rapid extinction begins, during which the population size decreases rapidly and selection is ineffective.

For the purpose of this paper, we reiterate three findings from these studies. Firstly, increasing the mutation rate has the strongest effect on reducing the time to extinction under mutational meltdown (41). This result is important because it indicates that the meltdown process can be induced by small alterations in the mutation rate (as attempted by mutagenic drugs) and that the resulting extinction is relatively robust to other parameters such as the carrying capacity (which is difficult to measure or control inside a host). Secondly, the variation in extinction times is relatively small for the same set of parameters (see Figure 1; 41). This result is important because it allows us to identify successful escape as an extension of the extinction time (both in simulations, and potentially also inside the host). Finally, the extinction time is minimized for intermediately strong deleterious selection coefficients (see Figure 1; 2). This is because Muller’s ratchet leads to the fastest loss in mean fitness when selection coefficients are too small to be purged immediately but large enough to reduce the mean fitness significantly (42). This result is important because it determines which types of deleterious mutations drive the meltdown process, and it is the basis for considering DFE modifiers as a potential rescue mechanism.

**Figure 1.**
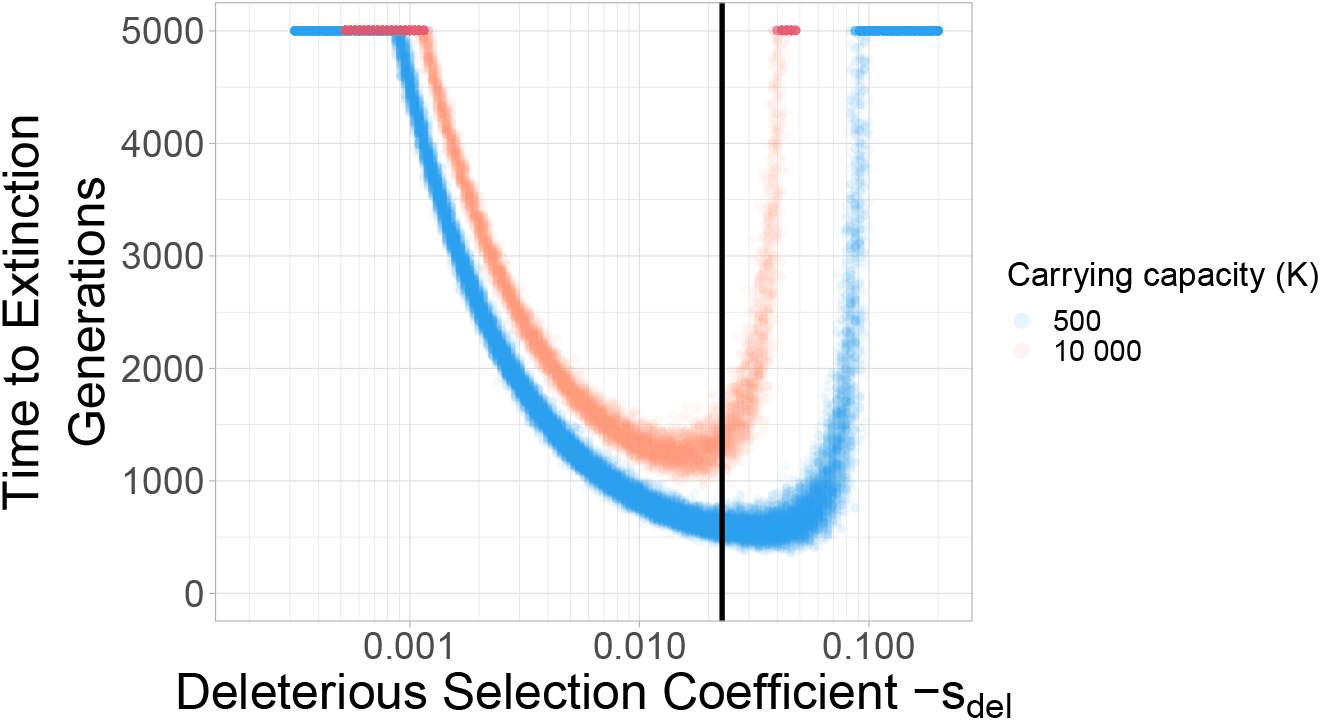
Deleterious selection coefficients of intermediate size minimize the extinction time. Two different carrying capacities are shown: 500 and 10 000. Each dot represents a simulation run. With larger carrying capacity *K* = 10000 the time to extinction increases. The black vertical line indicates the selection coefficient (*s*_del_ = *−* 0.023) used in the standard parameter set throughout the rest of the paper.

### 3.2 A continual input of growth-rate increasing mutations can halt Muller’s ratchet

Evolutionary theory often considers adaptation as a process of accumulation of beneficial mutations, which are mutations with a positive “selection coefficient”. Straightforwardly, these can be interpreted as growth-rate increasing mutations. For example, a resistance mutation often increases the growth rate of a pathogen in the presence of a drug. Here, we studied how beneficial mutations of equal size as the deleterious selection coefficient can help the population escape from mutational meltdown. Our simulations show that mutational meltdown can be prevented when a sufficient ratio of growth-rate increasing mutation is available. The ratio of beneficial to deleterious mutations that enables escape from mutational meltdown strongly depends on the carrying capacity of the population (see Figure 2). Here, the population can escape from mutational meltdown because Muller’s ratchet is halted by the continual input of beneficial mutations, as previously modeled by Silander et al. (43) and Goyal et al. (44).

**Figure 2.**
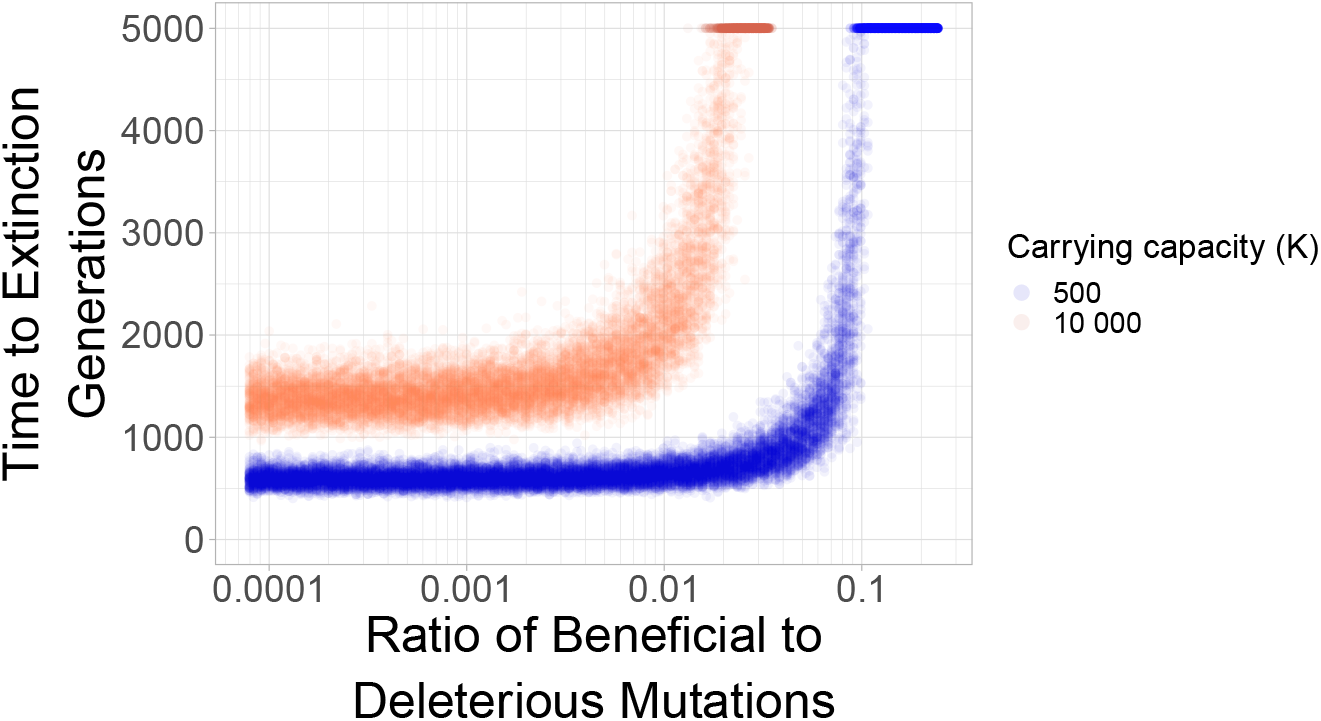
Mutational meltdown can be prevented with sufficient continual input of beneficial mutations. The sharp survival threshold depends strongly on the carrying capacity. The population can escape mutational meltdown when the ratio of beneficial to deleterious mutations is *≈*10%, when the carrying capacity is *K* = 500; a ratio of *≈*1% is sufficient for escape from mutational meltdown when *K* = 10000. Each dot represents a simulation run.

### 3.3 Reliable escape from mutational meltdown through mutation rate modifiers

Drug resistance occurs when a pathogen manages to “inactivate” the action of the drug such that it can normally propagate in the presence of the drug. In the case of a mutagenic drug, resistance occurs when a mutation alters the mutation rate of the virus in the presence of the drug, such that mutational meltdown is prevented. Here, we model such mutation rate modifiers, which occur at rate *U*_mutMod_ and which have effect size *f*_mutMod_. The occurrence rate *U*_mutMod_ determines the mutational target size (how frequently are such mutations available?) and the effect size indicates by how much the modifier mutation alters the mutation rate (i.e., *f*_mutMod_ = 0.1 means that the modifier decreases the mutation rate of its carriers to 10% of its previous value).

The effect size of the modifier strongly affects whether the population can escape mutational meltdown (Figure 3). The effect size of the modifier has to be large enough such that mutational meltdown is prevented at the modified mutation rate, had it been present from the beginning. Moreover, in order to allow for escape of the population from mutational meltdown, the resistance mutation has to appear early enough to be able to spread across the population before the lineage carrying the mutation modifier has accumulated too many deleterious mutations to escape extinction. Therefore, higher carrying capacities *K* and larger occurrence rates of the modifier *U*_mutMod_ facilitate resistance evolution by means of a mutation rate modifier. When the product of the carrying capacity and the occurrence rate of the modifier become small, whether the modifier invades and extends the extinction time becomes less deterministic, even for modifiers of large effect (Figure 3C).

**Figure 3.**
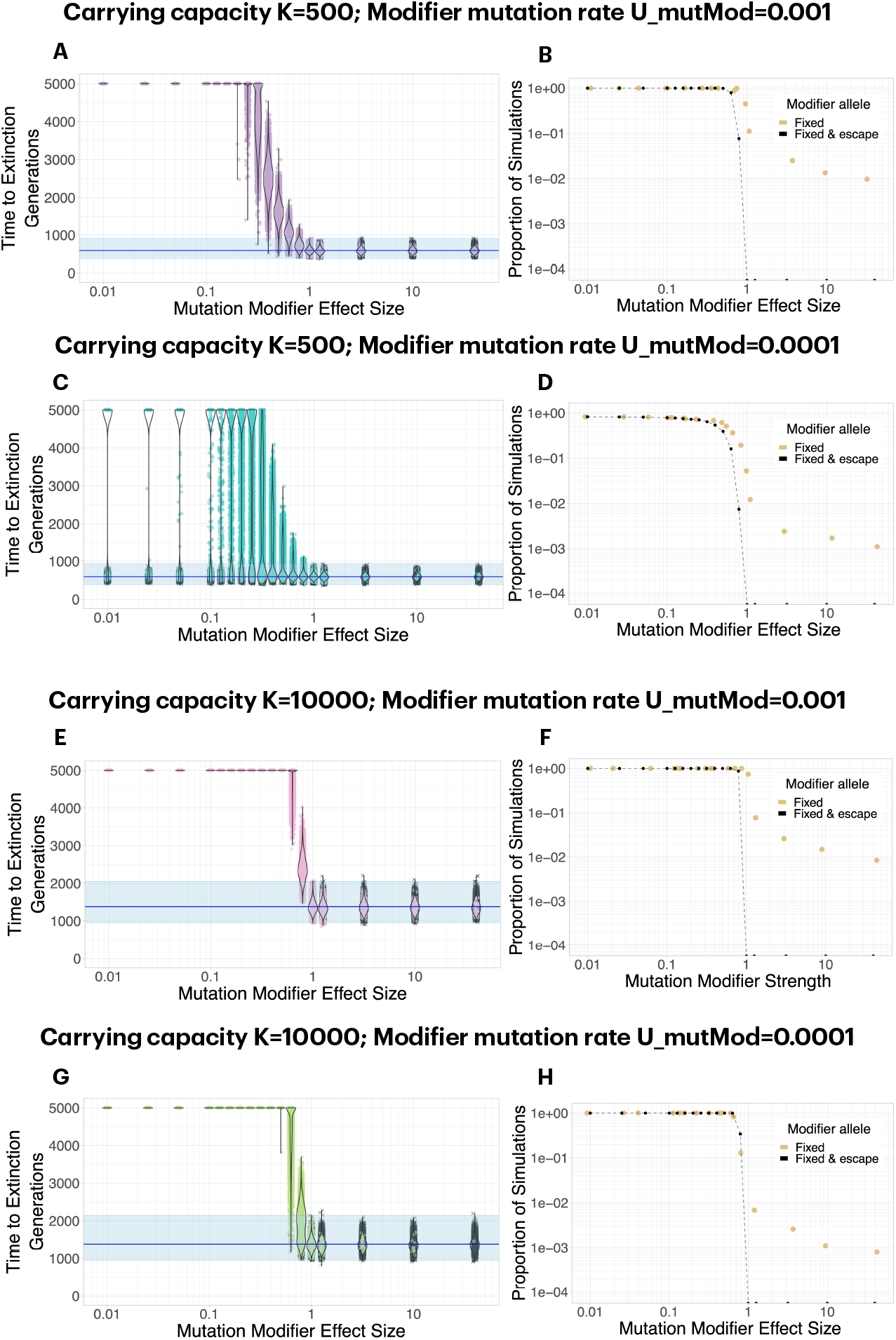
A mutation modifier of sufficient effect size can lead to escape from mutational meltdown. Each dot in Panels A, C, E, G represents a simulation run; simulations in which the mutation modifier was fixed in the population are shown in color, whereas simulations in which the modifier did not invade are shown in gray. Note that the effect size of the modifier, *f*_mutMod_, is defined multiplicatevely: e.g., if *f*_mutMod_ = 0.1, the mutation rate is reduced to 10% of its original value. B, D, F, H. In general, a mutation modifier that sufficiently decreases the mutation rate is effective at fixing in the population (yellow dots) and increasing the extinction time (black dots). Even when the modifier increases the mutation rate, it becomes fixed in some simulations, caused by random drift during the extinction phase (yellow dots for *f*_mutMod_ *>* 1). The graph shows the proportion of simulations for which the modifier allele was fixed (yellow dots), and the proportion of simulations for which it extended the extinction time beyond the maximum observed extinction time without the modifiers (black dots). C. For *K* = 500, escape from mutational meltdown becomes more stochastic with a lower modifier mutation rate. With *U*_mutMod_ = 0.0001, even for very strong modifiers, there is a small probability that escape is unsuccessful when the modifier allele appears too late. E-H. At higher carrying capacities *K* = 10000, escape from mutational meltdown is possible even with modifiers of relatively small effect, and the escape becomes more deterministic. Each parameter combination was simulated 10000 times.

### 3.4 Both negative and positive DFE modifiers can lead to escape from mutational meltdown

A modifier of the distribution of fitness effects can allow for escape from mutational meltdown in two ways. A DFE modifier can slow down Muller’s ratchet either by decreasing the effect of deleterious mutations (i.e., by weakening their effect) or by increasing their effect size (i.e., by making them more deleterious). This is true for intermediate-size deleterious mutations, for which the extinction time is smaller than for small or large-effect deleterious mutations (see Figure 1). We model DFE modifiers that occur at a rate *U*_dfeMod_ and with effect size *f*_dfeMod_. Again, the occurrence rate *U*_dfeMod_ determines the mutational target size. The effect size determines whether a DFE modifier is a “positive” modifier with 0 *< f*_dfeMod_ *<* 1, i.e., the effect of deleterious mutations is weakened, or whether it is a “negative” modifier with *f*_dfeMod_ *>* 1, i.e., the effect of deleterious mutations becomes stronger.

We find that both positive and negative DFE modifiers can lead to escape from mutational meltdown. The dynamics and probabilities of this escape differ greatly between the two types of modifiers (Figure 4, Supplementary Figure 1). For positive modifiers, a large effect size is necessary to lead to successful escape of the population from mutational meltdown for the whole duration of the simulations. When a positive DFE modifier has a sufficiently large effect size to lead to escape from mutational meltdown, it usually is successful at leading to escape from extinction, because all carriers of the modifier tend to be much fitter than the genotypes without the modifier. Thus, as soon as it appears, a positive DFE modifier provides a large benefit to its carriers. A negative DFE modifier acts fundamentally differently. It initially makes its carrier and their offspring less fit than any genotype with the same number of deleterious mutations that do not carry the DFE modifier. Thus, it initially has a (potentially) large deleterious effect on its carriers. Only when the subpopulation that carries the DFE modifier survives for several generations, the modifier becomes effective by means of efficient purging of new and segregating deleterious mutations. Thus, negative DFE modifiers rarely invade the population. When they invade, they lead to successful escape from mutational meltdown. We observe a strong dependence of the escape probability (and the invasion probability of a negative DFE modifier) on the product of the occurrence rate *U*_dfeMod_ and the carrying capacity *K*. That is because a negative DFE modifier has to appear on a genotype with no or very few deleterious mutations in order to be able to spread through the population, and this possibility is increased by maximizing the product of *U*_dfeMod_ and *K*.

**Figure 4.**
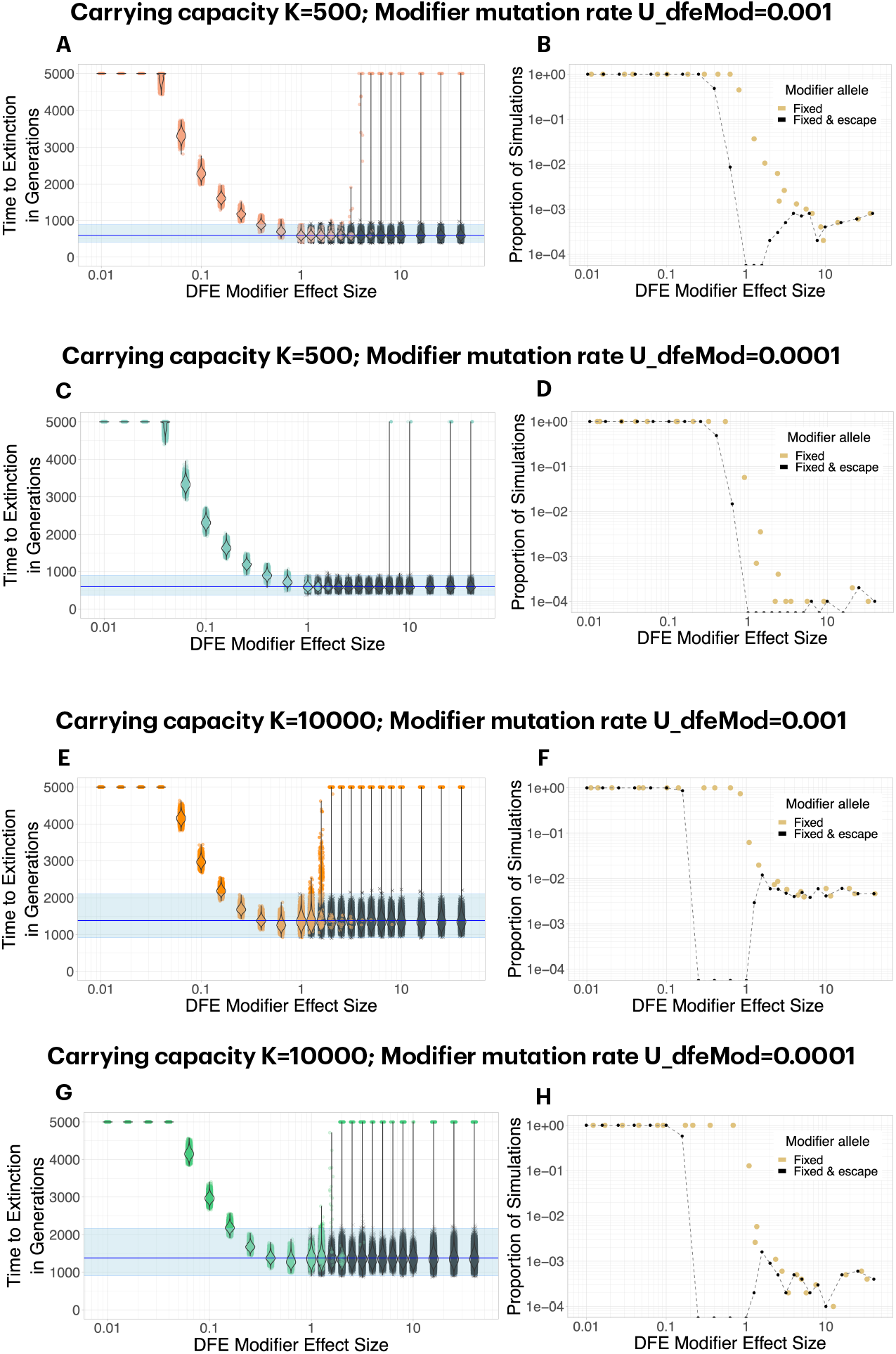
Both positive and negative DFE modifiers can lead to escape from mutational meltdown. Each dot in Panels A, C, E, G represents a simulation run; simulations in which the mutation modifier was fixed in the population are shown in color, whereas simulations in which the modifier did not invade are shown in gray. For positive DFE modifiers (effect size *f*_dfeMod_ *<* 1, i.e., the modifier makes deleterious mutations *less* deleterious), modifier alleles invade and fix reliably, but a large effect size *f*_dfeMod_ *<<* 1 is necessary to lead to successful escape from mutational meltdown (i.e., survival for 5000 generations). For negative DFE modifiers (effect size *f*_dfeMod_ *>* 1,, i.e., the modifier makes deleterious mutations *more* deleterious), modifier alleles invade and fix rarely, especially when the mutation rate to the modifier is low. Interestingly, although a negative DFE modifier initially tends to have a deleterious fitness effect on the lineage it carries, its escape probability increases with the carrying capacity (F, H), unlike what is expected for a deleterious mutation. Each parameter combination was simulated 10000 times.

## 4 DISCUSSION

Mutagenic drugs are under close investigation regarding potential treatment of RNA virus infections. Their mode of action is rooted deeply in evolutionary theory. Evolutionary theory also predicts mechanisms by which adaptation can occur. Here we propose and discuss three potential and fundamentally different adaptation mechanisms for adaptation to mutagenic drugs. We provide a proof-of-concept simulation study in order to encourage discussion about mechanisms of adaptation to mutagenic drugs and development of methods to infer such mechanisms. In addition, our work delivers a note of caution about potential rare-event adaptations that are almost impossible to predict with current experimental approaches.

### 4.1 Rethinking ‘beneficial’ mutations

The three adaptation mechanisms presented here act either by successively increasing the growth rate of the virus, by altering the mutation rate of the virus, or by altering the fitness effects of all deleterious mutations that the virus is accumulating (and has accumulated). Whereas all three adaptation mechanisms are ultimately beneficial to the virus population when they successfully lead to survival in the presence of the drug, only the first category consists of ‘beneficial’ mutations as they are usually considered in evolutionary biology.

Growth-rate increasing mutations as presented in the first model, do not provide resistance in the strict sense, as their continual input is necessary to allow for the survival of the population under high mutation rates over longer time scales. Notably, our model of growth-rate increasing mutations is greatly simplified by assuming that all beneficial mutations have the same effect size, which is the same effect size as that of deleterious mutations. Also, due to the neglect of back mutation in our model, beneficial (and deleterious) mutations occur at the same rate at all times. Alternative models of fitness increase that incorporate a fitness optimum, such as Fisher’s Geometric Model (see, e.g., 45), for example, are expected to yield different results regarding escape from mutational meltdown by means of growth-rate increasing mutations as compared to our simplified model. Largely independently of the underlying model, we expect that for successful escape, a serial compensation of the continually appearing deleterious mutations by beneficial mutations has to occur, whereas mutation and DFE modifiers can allow for a significant extension of the extinction time by means of a single mutation.

A mutation rate modifier that may reverse the mutation rate of the virus to its value in the absence of the drug can be considered a *drug resistance mutation*. That is because a population that carries a mutation rate modifier can proliferate in the presence of the mutagenic drug similar to the original genotype in the absence of the drug. Mutation-rate modifying mutations are frequently detected in experimental studies of microorganisms, for example in *E*.*coli* (e.g. 46, 47). However, both theoretical and empirical work are contentious as to how much selection acts on mutation rates and whether an equilibrium mutation rate will eventually be obtained (e.g., 48, 49, 50, 51, 52, 53, 54). Interestingly, the most well-studied adaptation to the mutagenic drug favipiravir, identified in the laboratory, was described as a mutation rate modifier (34).

Finally, a DFE modifier as defined here could be considered a *drug tolerance mutation*, because it does not alter the action of the drug, yet allows the virus population to survive in its presence. Here, the mutation rate of the virus remains high. At the same time, the altered selection coefficients of all mutations change the eco-evolutionary dynamics such that Muller’s ratchet becomes less severe and the population can cope with the increased mutation pressure. A DFE modifier is a specific form of epistasis (i.e., the genetic-background dependence of mutational effects), where the modifier allele alters mutational effects genome-wide. Sydykova et al. (55) have recently explored a related model of tunable epistasis, in which they found that epistasis can evolve in different directions under high mutation rates. Whether weakening the effects of deleterious mutations can lead to survival has been modeled in the research area of quasi-species theory, where a so-called “error threshold” determines the condition in which mutation overwhelms selection (e.g., for finite populations, 56, 57). This leads to the inability of a population to maintain its genetic information, which can in general be considered similar to mutational meltdown (58, but see 59). Moreover, the evolution of more severe effects of deleterious mutations has been theoretically studied with respect to models of drift robustness (60), quasi-species evolution (e.g., 61) and epidemiology (62).

There is a two-fold potential danger imposed by the successful invasion of a DFE modifier. Firstly, by changing the distribution of fitness effects of mutations globally, the presence of a DFE modifier may create a virus that evolves differently than its ancestral strain also in the absence of the drug, and hence in future infections of non-treated host. Secondly, by surviving in the treated host under high mutation rates, the virus population may be able to access adaptive genotypes in the sequence space that are not reachable under normal mutation rates (e.g., because they require multiple mutations). Although any prolonged survival of the virus population poses a danger regarding adaptation of the virus (to both the host immune system and the drug; 36), a DFE modifier is likely to facilitate such adaptation.

We are not aware of any empirical evidence of DFE-modifying adaptations in viruses. Of note, detecting such mutations is not straightforward because of their indirect effects, and a careful experimental design would be necessary to distinguish them from mutation rate modifiers, for example. Moreover, the different adaptation mechanisms described here could occur together (or interfere with each other), or many mutations of small DFE-changing-effect could be involved in adaptation to mutagenic drugs. In higher organisms, mutations in chaperones or global regulators could have the effect of DFE modifiers. For example, mutations of global transcription regulators, such as RpoB in *E. coli*, can change the expression patterns of hundreds or thousands of genes (63, 64). Chaperones (e.g., Hsp90 65 or DnaK 64) buffer the effects of mutations, and by weakening or strengthening the effect of the chaperone through mutation, the distribution of fitness effects would likely be altered either pathway- or genome-wide.

### 4.2 Negative DFE modifiers would not be detected in experimental screens for adaptations

An important result from the work presented here is that the escape probability for a very efficient adaptation mechanism (efficient if it invades the population) can be very small. Specifically, a negative DFE modifier, i.e., a mutation that makes other mutations more deleterious, has a very low probability to establish in the population in the first place. That is because it initially harms its carriers by reducing their fitness; only in the longer term the more efficient purging of deleterious mutations becomes an advantage. Their rare invasion implies that negative DFE modifiers would in general not be predicted by laboratory evolution experiments that are commonly used to screen for potential adaptations to new drugs (e.g. 12, 33, 34). If the probability of invasion of a negative DFE modifier is 1/10000, thousands of replicates of an evolution experiment would be necessary to observe it in the lab, whereas usually only *≈*3 replicates are performed in such experiments. However, once millions of patients are treated with the drug, the mutation would have ample opportunity to invade and spread. That is why theoretical and computational discussion and study of adaptation mechanisms are essential to complement experimental studies.

### 4.3 How to dissect the adaptation mechanism of a candidate mutation

Once candidate mutations of adaptation to mutagenic drugs have been identified (either from laboratory experiments or from the wild), it will be important to dissect the underlying adaptation mechanism. We propose that this could be possible by comparing the eco-evolutionary dynamics of virus population with and without the candidate mutation in the presence and absence of the drug. For example, if a resistance mutation reduces the mutation rate, less segregating and fixed mutations should be observed over time when comparing a population that carries this mutation with a non-mutated reference type. Moreover, if the resistance mutation alters the mutation rate independent of the drug treatment, the same pattern should be observed in non-treated populations. Interestingly, negative DFE modifiers should also display a pattern of reduced mutation accumulation in the presence and absence of the drug, because selection against deleterious mutations becomes more efficient. However, a DFE modifier does not prevent deleterious mutations from appearing, and instead leads to stronger selection against them after their appearance. We thus expect that the observed frequency spectrum of segregating mutations is different between a DFE modifier and a mutation rate modifier, with more low-frequency mutations for the DFE modifier. Conversely, since a positive DFE modifier reduces selection against deleterious mutations while leaving the mutation rate elevated, its signature should include elevated mutation accumulation compared to the reference type. Future work should further explore the predicted signatures of different adaptation mechanisms.

### 4.4 Limitations and future steps

The goal of this study was to conceptually introduce different types of adaptations to mutagenic drugs and highlight and discuss their different action from an evolutionary point of view. Our results should be seen as a qualitative illustration of potential effects of different adaptation mechanisms on looming extinction, each of which we study separately, but we caution against interpreting the reported numbers at face value. Our model is a simple depiction of intra-host evolution, in which a small number (*N*_0_ = 10) of mutation-free genotypes infect a host and grow exponentially to a predetermined carrying capacity (*K* = 500 or *K* = 10000) that is maintained until the mean growth rate of the population falls below 1. The increased mutation rate (as caused by a mutagenic drug) is present from generation zero onwards, whereas in natural scenarios treatment would only begin after the infection was detected. We consider as escape from mutational meltdown when a population that would have gone extinct without the adaptation mechanism in question survives for *t*_max_ = 5000 generations. In natural populations, *t*_max_ may reflect the time by which transmission has occurred, or by which the immune system is responding. Because we do not know the population dynamics of real virus populations in the wild, our parameters are somewhat arbitrarily chosen. For example regarding the initial population size, recent evidence has pointed towards infection bottlenecks (in influenza) being very small, with sometimes a single virion infecting a new host ((66, 67)). Moreover, although the census size of virions inside a host is probably on the order of millions, a much lower “effective” population size could reflect the evolutionary dynamics within a human host. In laboratory evolution experiments, the constant population size that best describes the evolutionary dynamics of populations under serial passaging is usually on the order of few hundreds, which is why we chose *K* = 500 as one of the studied population sizes (68, 33). Reassuringly, our results showed that the proposed adaptation mechanisms occurred at both tested population sizes, and especially for the DFE modifier (and surprisingly for both negative and positive DFE modifiers), we observed larger escape probabilities for larger population sizes, indicating that this escape mechanism is relevant also for (much) larger populations sizes (and thus, across all regimes of the dynamics of Muller’s ratchet, see, e.g. 69). We further ignore potential back mutations, and we do not consider the ability of some RNA viruses (including influenza) to rearrange, which is a type of recombination between genomic segments that could slow down Muller’s ratchet and stall extinction. Finally, we only consider a single, constant, deleterious effect that is common to every deleterious mutation as opposed to the distribution of fitness effects that is observed in natural and laboratory populations (e.g., 70). Relieving some of these assumptions is an important direction for future work.

### 4.5 Relevance for the use of mutagenic drugs in human populations

As argued above, we have studied adaptation to mutagenic drugs in a very simplified model that makes no attempt at reflecting the complexity of a human host. Nevertheless, our work cautions against the routine use of mutagenic drugs in human populations by demonstrating that there is much to learn about potential adaptation to mutagenic drugs. Adding to the arguments proposed by Nelson and Otto (36), our work shows that unpredictable types of adaptations can occur under wide-spread use of mutagenic drugs. Specifically, we consider tolerance mutations by means of a negative DFE modifier a type of adaptation that may affect evolution not only in current but also future (untreated) hosts. Moreover, the probability of adaptations (to both the drug and the host immune system) occurring and spreading to new hosts further increases when a mutagenic drug is not administered at sufficiently high doses or until the meltdown process is completed, as would be the case if a mutagenic drug is approved for household use.

## 5 CONTRIBUTION TO THE FIELD

Mutagenic drugs are currently discussed regarding their potential use against RNA viruses such as influenza and Sars-Cov-2. The principle of their action is rooted in the evolutionary theory of mutational meltdown, according to which the increasing load of deleterious mutations leads to collapse and extinction of a population. It is contentious whether and how RNA viruses can develop resistance to mutagenic drugs, and thereby escape mutational meltdown. Here, we present and discuss three potential mechanisms of adaptation to mutagenic drugs and model the eco-evolutionary dynamics when such mutations are available. Whereas growth-rate increasing mutations have to be continually available to prevent extinction of the virus population under drug treatment, modifiers of the mutation rate or the fitness effects of deleterious mutations can allow escape from mutational meltdown with a single mutation. We highlight the stochastic nature of this process that will rarely be captured by laboratory experiments. Moreover, we discuss the different adaptation mechanisms as examples of drug resistances versus drug tolerance.

## ACKNOWLEDGMENTS

The authors are grateful for discussion of the project in the THEE/Evoldynamics lab. Calculations were performed on UBELIX (http://www.id.unibe.ch/hpc), the HPC cluster at the University of Bern.

## DATA AVAILABILITY STATEMENT

The complete annotated documentation of the computational analyses of this paper will be archived on Zenodo upon publication of the paper and is currently deposited at https://gitlab.com/evoldynamics/ms-mutational-meltdown-2021.

