## Supplementary Figure 1 for "Evolutionary models predict potential mechanisms of escape from mutational meltdown"

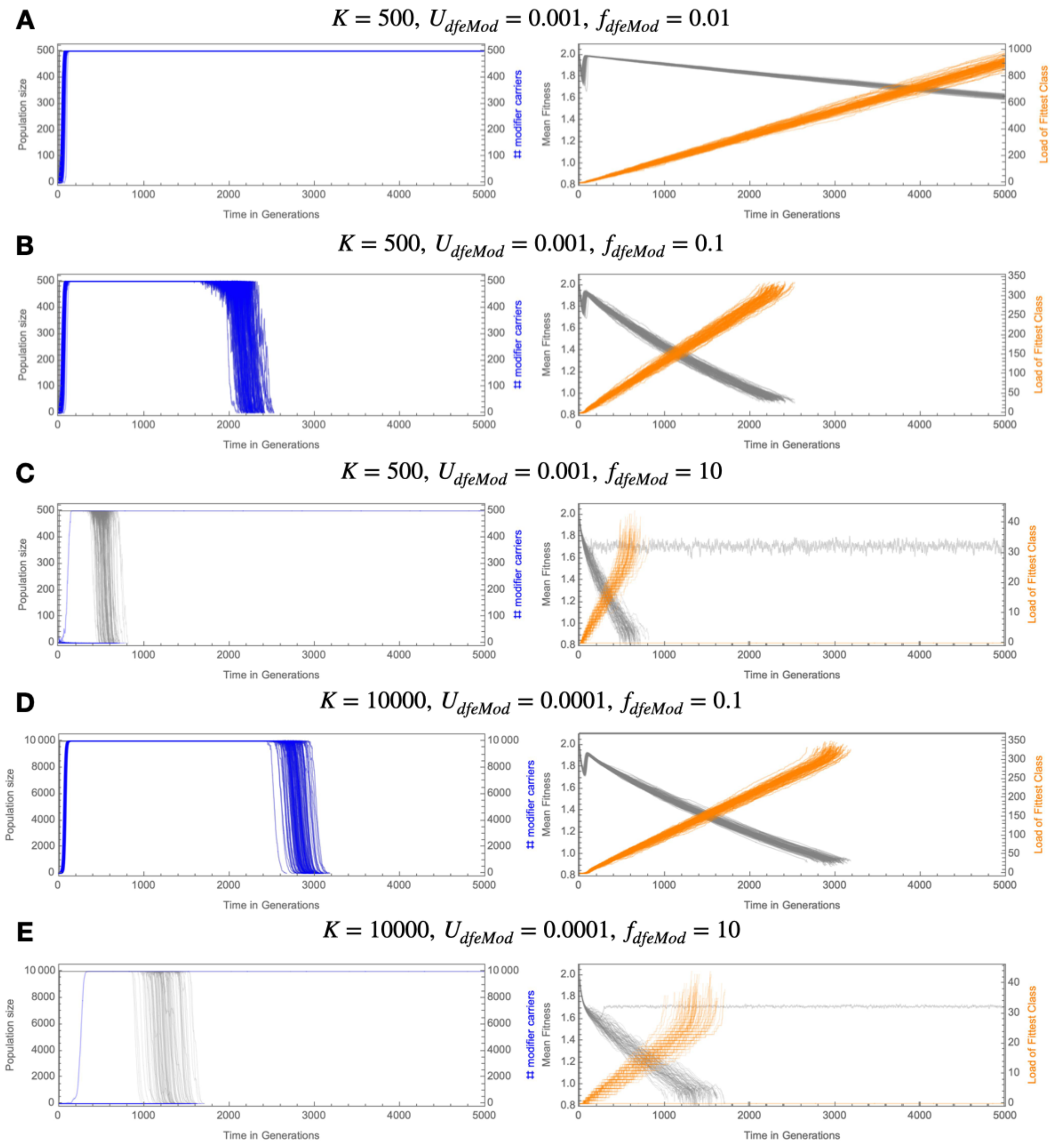


**Supplementary Figure 1.** Trajectories of population size (gray, left panels), number of carriers of the modifier (blue, left panels), mean population fitness (gray, right panels) and deleterious mutation load of the fittest genotype (orange, right panels) for five chosen parameter combinations in the DFE modifier model. 100 trajectories are shown per panel. A. For a strong positive DFE modifier, the modifier allele always invades and leads to survival for 5000 generations (left). Meanwhile, Muller's ratchet reduces the mean fitness and increases the mutation load in a linear fashion (right). B & D. For positive DFE modifiers of intermediate effect size, the modifier invades but does not extend the extinction time beyond 5000 generations. C & E. For a negative DFE modifier, invasion is very rare (here, only one instance of escape is shown per panel). If the modifier invades, it leads to survival of the population beyond 5000 generations.
